# ZSeeker: An optimized algorithm for Z-DNA detection in genomic sequences

**DOI:** 10.1101/2025.02.07.637205

**Authors:** Guliang Wang, Ioannis Mouratidis, Kimonas Provatas, Nikol Chantzi, Michail Patsakis, Ilias Georgakopoulos-Soares, Karen Vasquez

## Abstract

Z-DNA is an alternative left-handed helical form of DNA with a zigzag-shaped backbone that differs from the right-handed canonical B-DNA helix. Z-DNA has been implicated in various biological processes, including transcription, replication, and DNA repair, and can induce genetic instability. Repetitive sequences of alternating purines and pyrimidines have the potential to adopt Z-DNA structures. ZSeeker is a novel computational tool developed for the accurate detection of potential Z-DNA-forming sequences in genomes, addressing limitations of prior methods. By introducing a novel methodology informed and validated by experimental data, ZSeeker enables the refined detection of potential Z-DNA-forming sequences. Built both as a standalone Python package and as an accessible web interface, ZSeeker allows users to input genomic sequences, adjust detection parameters, and view potential Z-DNA sequence distributions and Z-scores via downloadable visualizations. Our Web Platform provides a no-code solution for Z-DNA identification, with a focus on accessibility, user-friendliness, speed and customizability. By providing efficient, high-throughput analysis and enhanced detection accuracy, ZSeeker has the potential to support significant advancements in understanding the roles of Z-DNA in normal cellular functions, genetic instability, and its implications in human diseases.

**Availability:** ZSeeker is released as a Python package under the GPL license as a multi-platform application and is available at: https://github.com/Georgakopoulos-Soares-lab/ZSeeker. A web-interface of ZSeeker is publicly available at https://zseeker.netlify.app/.

## Introduction

Since the discovery of the canonical right-handed B-DNA double helical structure, a number of alternative DNA conformations (i.e. non-B DNA) that differ from the canonical “Watson-Crick” B-form DNA structure (Watson and Crick 1953) (e.g. Z-DNA, H-DNA, G4-DNA, and hairpin/cruciform DNA) have been characterized (G. Wang and Vasquez 2022; Herbert 2020; Kasinathan and Henikoff 2018). Z-DNA, a left-handed DNA structure (A. H. Wang et al. 1979), plays important roles in various genomic functions and has profound implications for transcription, replication, nucleosome positioning, recombination, DNA damage and repair (Zavarykina, Atkarskaya, and Zhizhina 2019; G. Wang and Vasquez 2007; Georgakopoulos-Soares, Chan, et al. 2022; Wong et al. 2007). Z-DNA-forming sequences have been shown to stimulate genetic instability and are enriched at mutation hotspots in human cancer genomes, implicating them in the etiology of human diseases (McKinney et al. 2020; G. Wang, Christensen, and Vasquez 2006; Georgakopoulos-Soares et al. 2018; Bacolla et al. 2016; Lu et al. 2015; Bacolla et al. 2019).

Alternating purine-pyrimidine sequences have the capacity to adopt Z-DNA structures, in which the purine residues are in the syn-conformation and pyrimidines in the anti-conformation. The sugar-phosphate backbone of the DNA helix is twisted into a left-handed zigzag-shape by the alternating syn- and anti-conformations (Rich and Zhang 2003). However, it has been realized that not all alternating purine-pyrimidine dinucleotides have thermodynamic features that promote Z-DNA formation, with GC being the most favored, followed by GT/AC, and AT the least stable (Jovin et al. 1983; Peck and Wang 1983).

Based on the unique sequence features required for Z-DNA formation, multiple sequence-based computer algorithms, evaluating sequence type and length were developed to search for potential Z-DNA-forming sequences in genomes (Ho et al. 1986; Cer et al. 2010). The energetic parameters for stabilizing all sixteen possible dinucleotides in the Z-DNA conformation were calculated (Ho et al. 1986) and this information was used to develop the well-recognized “Z-Hunt” program to predict potential Z-DNA-forming sequences in genomes (Peck and Wang 1983).

The development of the Z-Hunt program was a major step forward for computer aided DNA structure prediction and was recognized as the most accepted model because it was based on the energetic requirements for Z-DNA formation estimated from experimental data. However, certain limitations of these foundational studies should be noted. For example, it is known that divalent cations such as magnesium (Mg^2^□), calcium (Ca^2^□) and zinc (Zn^2^□) can stabilize Z-DNA structures, while low concentrations of alkali cations such as sodium (Na□) and potassium (K□) favor the B-DNA conformation. This was not considered in the energetic measurements of Z-DNA formation *in vitro* that were used for the algorithm design. In addition, these experiments were performed on plasmid DNA used for these early studies containing multiple non-B DNA-forming motifs in addition to Z-DNA that could compete with each other in the structure transition measurements, confounding the interpretation of the Z-DNA-forming potential. Further, the algorithm was developed based on knowledge and technology several decades ago, and thus has several limitations for the accurate prediction of Z-DNA-forming sequences. The NCI non-B DB (https://nonb-abcc.ncifcrf.gov/apps/site/resources) Genomic Database Search Tool also uses Z-Hunt principles for Z-DNA searching and scoring, but gives significantly different predictions to Z-Hunt, as discussed later in our comparison section (Cer et al. 2010). This current state underscores the need for an accurate, easy-to-use, and reliable tool to predict Z-DNA structure formation which can help catalyze further research in the field.

Limited by the computer processing ability at the time of its development, the Z-Hunt algorithm searches long sequences by subdividing the sequences into fixed search windows of 16-24 bps and evaluating the Z-DNA-forming potential in each window. Although the windows have overlap to reduce the possibility of dissecting Z-DNA-forming motifs between two adjacent windows, it natively has limitations in identifying long Z-DNA-forming sequences, particularly those that have sequence interruptions but still maintain Z-DNA-forming capabilities.

Z-Hunt examines each window independently, calculating the energies for Z-DNA formation at the dinucleotide level and does not account for potential longer-range effects, such as DNA bending forces (Yi, Yeou, and Lee 2022). However, the energy penalties for forming a Z-DNA structure are more complicated than adding the energy for each dinucleotide. For example, the unpaired and extruded bases at the B-Z junctions allow the adjoining Z-DNA segment to base stack on the neighboring B-DNA segment, which requires more energy than maintaining the alternating purine/pyrimidine dinucleotide in the Z-DNA formation (Zhabinskaya and Benham 2011). Therefore, the average free energies for each dinucleotide in a longer Z-DNA-forming motif may be less than the same dinucleotide in a short motif which also must flip two B-Z junctions. However, when two Z-DNA motifs with scores X and Y were put together, the merged sequence received a combined score of X+Y in Z-Hunt, even though the merging reduces two out of four energy-consuming B-Z junctions.

Recently, deep learning approaches were implemented to search for potential Z-DNA-forming sequences (Beknazarov, Jin, and Poptsova 2020). The accuracy of deep learning approaches depends on the availability of high-quality large datasets for training, and is convoluted by ADAR1A (a Z-DNA-binding protein used in the ChIP-seq to develop the programs) binding preferences (Marshall et al. 2020).

Building on the experimental thermodynamic data generated previously, here we also included data from biochemical and genetic experiments of Z-DNA formation from different systems, along with advanced computational processing ability, to design a novel and optimized Z-DNA-finder program. This program allows for a more accurate and flexible Z-DNA predicting and searching platform than the currently existing programs. We introduce “ZSeeker”, a Python package that enables rapid and accurate identification of Z-DNA-forming sequences. We also provide a web-portal for analyzing sequences for Z-DNA structure-forming potential, visualizing and downloading the outputs (**Figure 1**). With ZSeeker, we aim to support the scientific community in advancing Z-DNA research and its applications.

**Figure 1:**
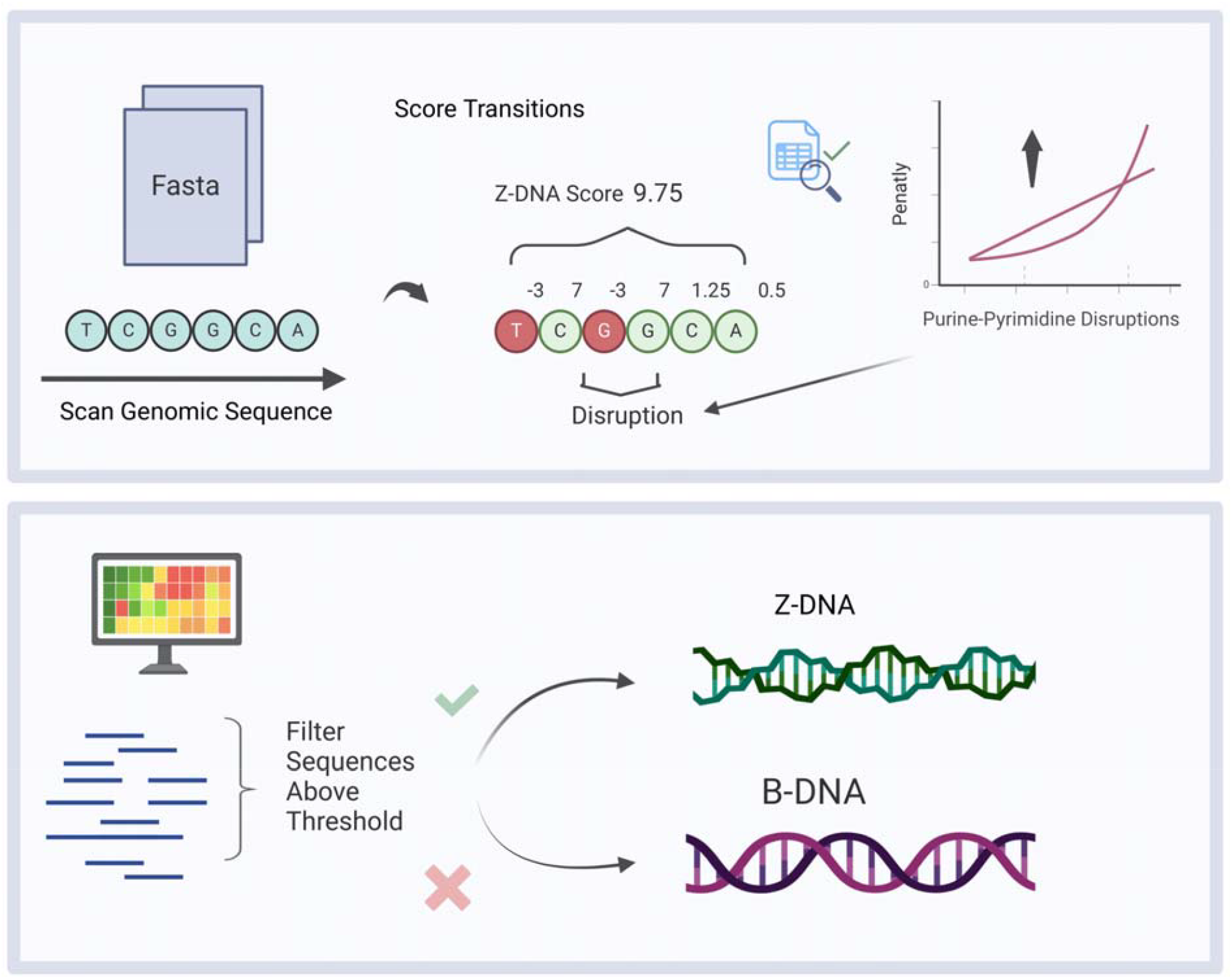
Schematic representation of B-DNA and Z-DNA conformations as well as the algorithmic design of ZSeeker. The schematic also illustrates the scoring algorithm and penalties that ZSeeker allows.

## Results

### Prediction parameters

Since the energetic parameters from biochemical experiments do not fully mimic the complex conditions *in vivo*, and detecting Z-DNA formation directly in living cells is challenging, we used Z-DNA-induced mutagenesis as an indirect indicator for Z-DNA formation *in viv*o. We found that a CG(5) repeat cloned into the lacZ’ gene in a mutation reporter was ∼30x more mutagenic than control B-DNA (puCON) in bacterial DH5alpha cells; and when the CG repeats were increased to 14 units CG(14), the mutation frequencies increased further in bacteria, mammalian cells, and in mice (G. Wang, Christensen, and Vasquez 2006; G. Wang et al. 2008). S1 enzyme probing and chloroquine gel electrophoresis analysis also confirmed the Z-DNA conformation in the CG(14) repeat. Under the same conditions, a GT(14) repeat was not mutagenic (**Figure S1**), and a GT20 repeat cloned in a *supF* mutation reporter only induced slightly higher mutation frequencies than the control B-DNA sequence in mammalian COS-7 cells. In addition, most of the mutants were small in/dels within the repeats (**Figure S2**), likely due to slipped strand mispairing occuring on simple repeats (Kelkar et al. 2010), rather than the signature Z-DNA-induced large deletions resulting from DNA double-strand breaks (DSBs) as seen in mammalian cells (G. Wang, Christensen, and Vasquez 2006), suggesting a lack of stable Z-DNA formation *in vivo*. When the GT repeat reached a length of 30 (GT30), although only slightly more mutagenic than AT14 and GT20, it stimulated deletions in >50% of the mutants, a signature Z-DNA-induced mutation in mammalian cells. A longer GT41 repeat induced similar levels of mutation in COS-7 cells as a CG14 sequence, and the majority of mutants were large deletions resulting from the formation of DSBs, providing strong evidence for Z-DNA formation in mammalian cells (Xie et al. 2019). In summary, our *in vivo* experiments suggested more substantial differences between the GC and GT repeats where GC sequences only required 5 repeats to form a Z-DNA structure (**Figure S1**), while GT sequences required 30 repeats to adopt a Z-DNA structure (**Figure S2**), consistent with previous reports (A. H. Wang et al. 1984; Casasnovas et al. 1989; Kim, Yang, and DasSarma 1996). For AT dinucleotides, we found that a GC repeat containing 2 embedded AT repeats can still form Z-DNA (G. Wang and Vasquez 2006). However, it was found that an AT4 repeat, even when located between two CG6 repeats, adopted an unwound structure rather than a Z-DNA structure (Ellison et al. 1986), and an AT14 repeat in a *supF* reporter adopted slippage bubble structures instead of Z-DNA, resulting in small indels within the repeats rather than DSBs and large deletions (**Figure S2**). Based on historical and current experimental data, we designed our new Zseeker prediction and searching program.

## Materials and Methods

The succession of bases within repetitive sequences dictates the potential interactions among different regions in a DNA molecule and determines its potential for the formation and stability of non-B DNA structures. Although the formation of non-B DNA is affected by many other factors such as salt, pH, negative supercoiling, and the presence of binding proteins, the linear sequence feature is the premise for the ability of a sequence to adopt secondary structural conformations. We provide below the methodology and parameters of our sequence-based computer algorithm designed to search sequences uploaded by the user for repeat tracts with the propensity to adopt Z-DNA structures.

### Algorithmic process for Z-DNA detection

An alternating purine-pyrimidine sequence such as a GC or GT repeat has the propensity to adopt a Z-DNA structure, in which purine residues are in the syn-conformation and pyrimidines are in the anti-conformation. The alternating syn- and anti-formations twist the sugar-phosphate backbone into a left-handed zigzag-shape (Rich and Zhang 2003). Under certain conditions such as base modifications or the presence of ions, Z-DNA structures can also form on regions that are not typical alternating purine-pyrimidine patterns, such as CCG/CGG or CGATCG repeats (Latha et al. 2002; A. H. Wang et al. 1985). Results from our biochemical and genetic experiments suggested that on a ∼7,000-bp plasmid (pMB1 replication origin) prepared from DH5alpha bacterial cells, a GC repeat could form a Z-DNA structure when it contained >4 GC repeats; GT repeats could form a Z-DNA structure when the sequence contained >25 repeats; while an AT repeat preferentially formed hairpins or loops rather than Z-DNA.

This algorithm scores the potential Z-DNA-forming sequences based on the transitions between the bps that support or prevent Z-DNA formation. Since a GC(4) repeat has 7 “G to C transitions”, and a GT(25) repeat has 49 “G to T transitions”, and both are minimum lengths for Z-DNA conformation, this algorithm gives each G to C or C to G transition a score of “7”, each “G to T” or “T to G” transition a score of “1”, and sets the threshold for Z-DNA formation at “50”. An “A to T” or “T to A” transition scores 0.5 (**Table 1**). However, a continuous AT or TA run over 4 repeat units is more likely to form hairpins or cruciform structures, and therefore has a high penalty score (**Table 1**). A sequence obtaining a score over the threshold will be considered as a candidate sequence. The algorithm is designed to identify the DNA subsequences that achieve the highest score and mark them as potential Z-DNA-forming sequences.

The recommended parameters for Z-DNA searching are provided below in Table 1:

**Table 1:** Recommended parameters for Z-DNA searching.

| Parameter | Description | Value |
| --- | --- | --- |
| Threshold | Scoring threshold for a sequence to be considered a potential Z-DNA-forming sequence and returned by the program | 50 |
| GC weight | Weight given to GC and CG transitions | 7 |
| AT weight | Weight given to AT and TA transitions | 0.5 |
| Consecutive AT scoring* | Penalty array for consecutive AT/TA repeats to account for hairpin structures. Provides the score adjustment for each AT/TA transition | 0.5, 0.5, 0.5, 0, 0, -5, -100 |
| AC weight | Weight given to AC and CA transitions | 1.25 |
| GT weight | Weight given to GT and TG transitions | 1.25 |
| Mismatch penalty starting value** | Penalty applied to the first non-purine/pyrimidine transition encountered | 3 |
| Mismatch penalty type | Method of scaling the penalty for contiguous non-purine/pyrimidine transitions | Linear or Exponential |
| Mismatch penalty linear delta*** | Rate of increase of the penalty for each subsequent non-purine/pyrimidine transition (only applies if penalty type is linear) | 3 |

**\*Consecutive AT scoring:** Because over 4 consecutive AT/TA repeats tend to form hairpin structures instead of Z-DNA (Ellison et al. 1986), a penalty array is defined, which provides the score adjustment for the first and the subsequent AT/TA transitions. The last element will be applied to every subsequent AT/TA repeat.
**\*\*Mismatch penalty starting value:** Penalty applied to the first non purine/pyrimidine transition encountered: 3.
**\*\*\*Mismatch penalty linear delta:** Only applies if penalty type is set to linear. Determines the rate of increase of the penalty for every subsequent non purine/pyrimidine transition: 3
NOTE: Non-B DNA formation is a dynamic process and is not decided solely by the DNA sequence. Cellular activities such as replication, transcription, and DNA repair can impact non-B DNA formation via changing DNA supercoiling levels, affecting DNA binding proteins, and modulating nucleosome/chromatin structures. The results from this algorithm only provide initial information on the potential of a given DNA sequence to form a Z-DNA structure.

### CLI Tool and Package

ZSeeker is available both as a command-line interface tool and as a Python library, distributed freely as an open-source package on PyPI (https://pypi.org/project/ZSeeker/). Users can install ZSeeker with a single command and choose to use it through the command line or integrate it into their Python code. Both options allow users to adjust the tool’s performance by utilizing multiprocessing to evaluate input sequences more efficiently.

### ZSeeker Web-interface

The ZSeeker web interface provides a straightforward, accessible version of the ZSeeker command-line tool. It uses the Gin framework in Go for the backend and HTML, CSS, and JavaScript for the frontend. The main purpose of the web application is to make ZSeeker’s features easy to use. Users can submit jobs, inspect, download, and filter Z-DNA results directly through the interface. It also includes visualizations to help users identify and evaluate potential Z-DNA regions. Additionally, there is an About/Contact page with information on the tool’s algorithms and a help page explaining its features and functions.

The top navigation bar is composed of three tabs, the “Job Submission”, the “About/Contact” and “Help” tabs, enabling navigation across the different parts of the ZSeeker website (**Figure 2A**). The “Job Submission” page displays default parameters for performing Z-DNA detection, which can be adjusted by the user for their specific needs. The user can either paste a FASTA sequence in the input box or upload a FASTA file (**Figure 2B**). After pressing the submit button, the file is processed, returning a set of outputs. These include a table and a CSV file with all potential Z-DNA sequences detected (**Figure 2C**), and two visualizations, a scatter plot showing the Z-score and Z-DNA sequence length for all sequences detected, and a boxplot displaying the distribution of Z-scores for the input data (**Figure 2D**).

**Figure 2:**
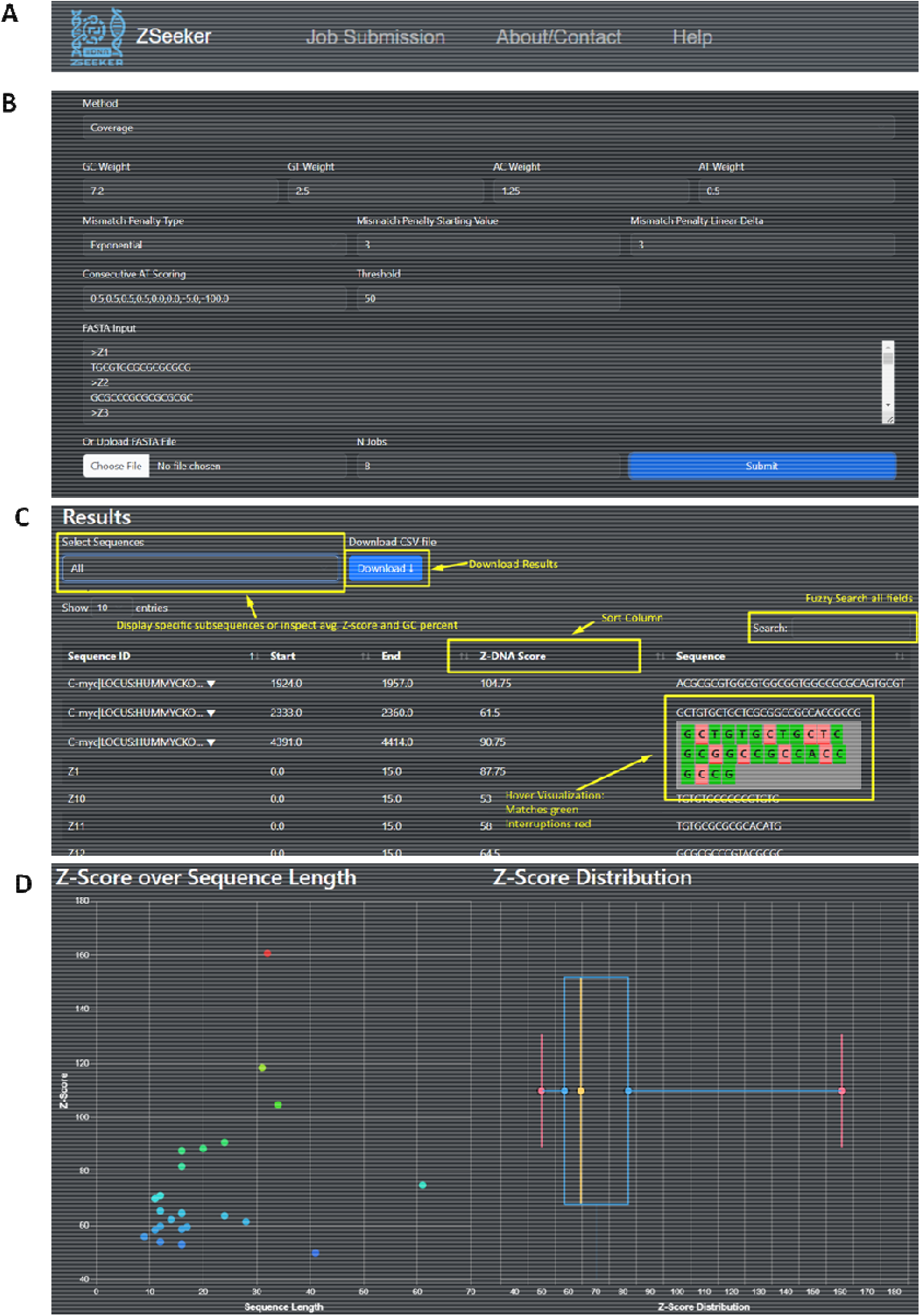
ZSeeker front-page explaining the navigation to different pages and features. **A**. Navigation bar for ZSeeker front-pages. **B**. The Job Submission page enables algorithm fine tuning. **C**. The results section on the Job Submission page shows the Z-DNA sequences above the threshold score and provides users with the option filter, download, and sort features. **D**. Visualizations are used to gain a quick overview of the results dataset.

The website contains a Help page, which provides information about Z-DNA to introduce the users to the ZSeeker program and guidance to the different features of the web app (**Figure 3A**). An About/Contact page provides contact information for potential bug fixes and feature requests (**Figure 3B**). A Privacy and License page is integrated that includes the website’s security, policies on personal data collection (**Figure 3C**) and the license used, which is Creative Commons 4.0 BY-SA (**Figure 3D**).

**Figure 3:**
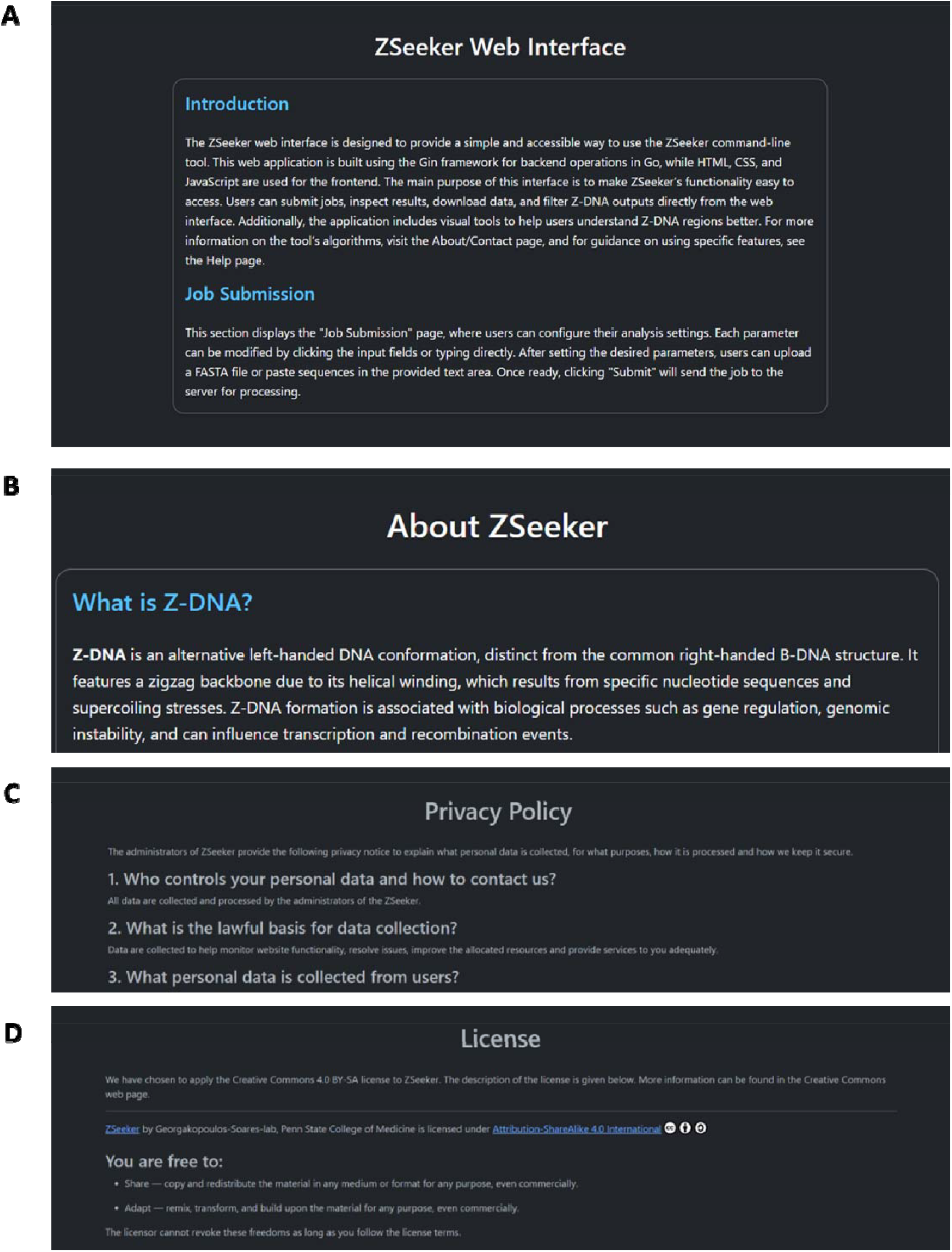
ZSeeker Help, About, and Privacy policy & License pages. A. Help Page, **B**. About Page, **C**. Privacy policy section **D**. License section.

### The accuracy and efficiency of ZSeeker compared to other available programs

Among the various Z-DNA prediction algorithms available, Z-Hunt and its improved version, Z-Hunt II, are notable for their foundation in thermodynamic parameters. These algorithms predict the formation of Z-DNA by analyzing the transition of all sixteen possible dinucleotides using a 2-D gel electrophoresis approach on negatively supercoiled plasmids in TBE buffer. As a result, Z-Hunt and Z-Hunt II have become widely recognized and accepted tools in the field. We tested the Z-Hunt II implementation currently provided by the Pui Shing Ho Lab (https://github.com/Ho-Lab-Colostate/zhunt) (Ho et al. 1986). As the online version of this tool (http://zhunt.bmb.colostate.edu/) was unavailable at the time of writing of this paper (June to November 2024), we used the offline version provided on GitHub. However, even with modern computational resources, this 40 year-old algorithm is extremely slow, with suboptimal architecture and lack of documentation, and it is cumbersome for the users. For example, we have encountered several limitations of the tool. Firstly, the algorithm has a time complexity of O(2^n^) due to its exhaustive search through all possible anti-syn conformations for a DNA sequence of length n dinucleotides. For instance, with a window size of 20 dinucleotides, the program must evaluate *2*^20^ (over one million) possible conformations for just one window, leading to extremely long computation times. This exponential growth makes the tool challenging to use for even medium-sized inputs unless the parameters minimum and maximum window sizes are set to very small values. Secondly, while our modified version outputs the nucleotide sequences, the original tool does not provide sequence coordinates within the larger DNA context and only includes the anti-syn structure with the lowest energy. This limitation reduces the practical utility of the results for biological applications where positional information is crucial. Additionally, the scores generated by the program, delta linking number, slope, and probability, are not clearly explained. There is insufficient documentation detailing what each score represents or which one should be considered indicative of Z-DNA propensity. Furthermore, there is no cutoff in the output results. The program includes all sequences in the output regardless of their scores, making it difficult to distinguish between sequences with a high potential to form Z-DNA and those with low potential. This absence of a threshold complicates the analysis and makes it challenging to focus on the most relevant sequences.

Additionally, we encountered limitations regarding sequence lengths. Due to the dinucleotide basis of the Z-Hunt II algorithm, it cannot directly score sequences of odd length as a whole. For example, when analyzing the seventeen-nucleotide sequence GCGCGTGCATATGCGTG, Z-Hunt II cannot evaluate the entire sequence because it requires sequences composed of an even number of nucleotides to form complete dinucleotides. This means the last nucleotide in an odd-length sequence cannot be incorporated into the analysis, potentially missing critical information. As a result, sequences such as GCGCGTGCATATGCGTG cannot be fully assessed by Z-Hunt II, limiting its applicability when dealing with sequences of varying lengths commonly found in genomic data. To demonstrate several of these challenges with Z-Hunt II, including its computational inefficiency, incomplete output, and lack of clear documentation, we used 20 Z-DNA-forming sequences that obtained Z-scores ranging from 31.7 to 1,750 in the original publication (Schroth, Chou, and Ho 1992) for this program, non-B_gfa and our novel algorithm, and the results are summarized in Table 2.

**Table 2:**
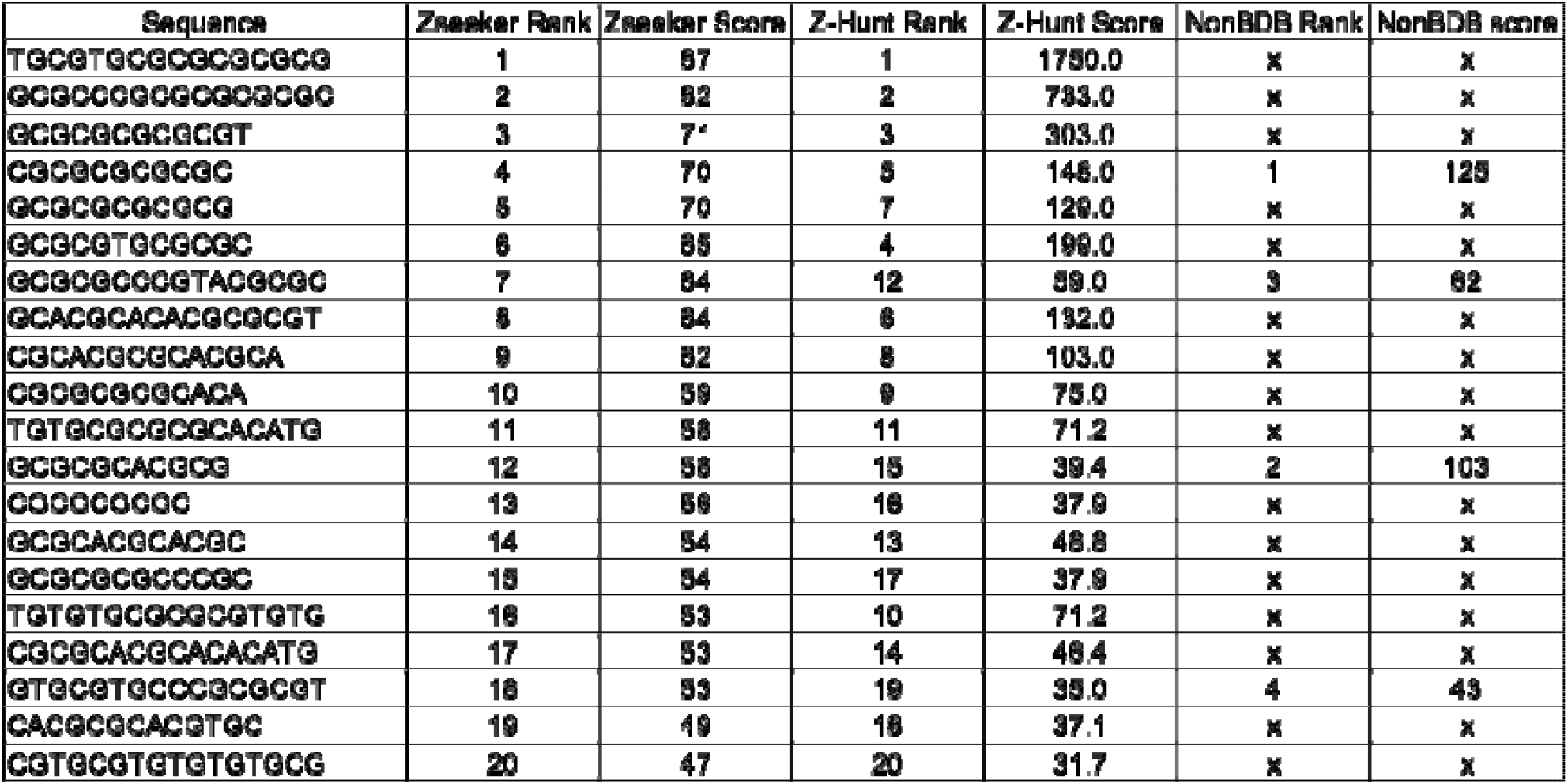
Comparing the Z-DNA scores from Z-Hunt II, NonBDB and Zseeker.

The non-B DNA Motif Search Tool (nBMST) in the NCI non-B DB Genomic Database (https://nonb-abcc.ncifcrf.gov/apps/site/resources) is a more user-friendly tool and is also very well accepted. Hence, we compared the search results from these two Z-DNA searching/prediction tools to the report from our ZSeeker program. Because Z-Hunt II (https://github.com/Ho-Lab-Colostate/zhunt) provides its output in an anti-syn structural form, we developed a modified version that logs instead the nucleotide sequences in the output to be able to compare with Zseeker’s results.

Despite their fundamentally different approaches, Z-Hunt and ZSeeker ranked the 20 sequences very similarly, as shown in Table 3. Notably, sequences ranked #7 and #16 by Zseeker were ranked #12 and #10 by Z-Hunt, respectively. In contrast, the non-B DNA Motif Search Tool (nBMST) failed to identify most (16/20) of the Z-DNA motifs that were reported in both ZSeeker and Z-Hunt, including those having the highest scores.

**Table 3:**
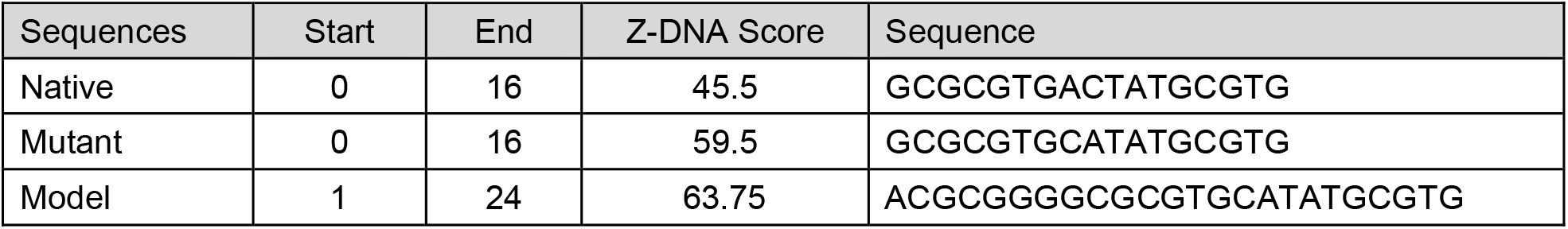
ZSeeker predictions.

To further compare the accuracies of ZSeeker and Z-Hunt in predicting Z-DNA-forming sequences, we tested sequences that have been confirmed to form Z-DNA structures in published literature using both programs. Visentin & Harley studied a sequence GCGCGTGACTATGCGTG in the mouse metallothionein I promoter (Visentin and Harley 1987) offering an ideal series of sample sequences to evaluate predictive accuracy. The native sequence has an alternating purine/pyrimidine element with an AC dinucleotide interruption in the middle. Reversing the interruption AC to CA created a consecutive alternating purine/pyrimidine pattern without changing the base contents. Two-dimensional chloroquine gel analysis and diethyl pyrocarbonate (DEP) sensitivity assays confirmed that a Z-DNA structure was formed on the mutant GCGCGTGCATATGCGTG sequence, while the native GCGCGTGACTATGCGTG was not able to adopt a Z-DNA structure unless a strong torsional stress was applied, while the CG(n) repeats and the GACGCGGGGCGCGTGCATATGCGTGG sequence formed Z-DNA structures even more readily than the mutant sequence (Visentin and Harley 1987). ZSeeker scored the native GCGCGTGACTATGCGTG 45.5, right below the 50 cut-off score; the mutant GCGCGTGCATATGCGTG was score 59.5 and was recognized as a Z-DNA-forming sequence; and the GACGCGGGGCGCGTGCATATGCGTGG sequence was cored 63.75 (Table 3). Thus, the predicting result matched the experimental results.

## Discussion

The development of ZSeeker represents a significant advance in the identification of sequences that have the potential to adopt Z-DNA structures. While previous algorithms such as Z-Hunt II laid foundational work in identifying potential Z-DNA regions, ZSeeker addresses significant gaps in computational efficiency, and by incorporating sequence imperfections, allows for more nuanced recognition of Z-DNA motifs. ZSeeker is efficient in predicting Z-DNA formation across genomic sequences. The results corroborate previous thermodynamic, biochemical, and cellular-based experimental data, indicating that GC repeats more readily form Z-DNA than GT or AT repeats.

Examination of Z-DNA-forming sequences with ZSeeker can be incorporated in studies related to genetic instability, genome organization and evolution, gene regulation, and human health and disease (Zhao et al. 2009; G. Wang and Vasquez 2022; Georgakopoulos-Soares, Victorino, et al. 2022; G. Wang and Vasquez 2007, 2006; Herbert 2019). ZSeeker features an intuitive web interface for seamless Z-DNA detection, offering outputs in the form of tables and visualizations that present Z-scores, penalties, and rank-ordered input sequences. Even though we recommend specific parameters to run Zseeker as described herein, parameters can be altered enabling fine-tuning of the ZSeeker parameters for personalized searches. For larger datasets and programmatic use, a Python package is also available. By providing more accurate and adaptable Z-DNA detection, ZSeeker addresses a critical gap in current genomic analysis tools. We anticipate that our novel Z-DNA detection tool will encourage further research and exploration of Z-DNA within the scientific community.

## Code Availability

The Zseeker package and its Python bindings are released under GPL license as a multi-platform application and are available at: https://github.com/Georgakopoulos-Soares-lab/ZSeeker

## Key Points

1. Z-DNA plays important roles in many cellular processes yet current search tools have limitations that we have addressed by developing a novel Z-DNA search tool
2. Zseeker is a novel computational tool developed for the accurate detection of potential Z-DNA-forming sequences based on experimental data
3. Zseeker functions both as a standalone Python package and as an accessible user-friendly and high throughput web interface

## Acknowledgements

This work was supported by grants from NIH/NCI R01CA093729 (to KMV) and from NIGMS under award number R35GM155468 (to IGS).

