## Supplementary Material for "ZSeeker: An optimized algorithm for Z-DNA detection in genomic sequences"

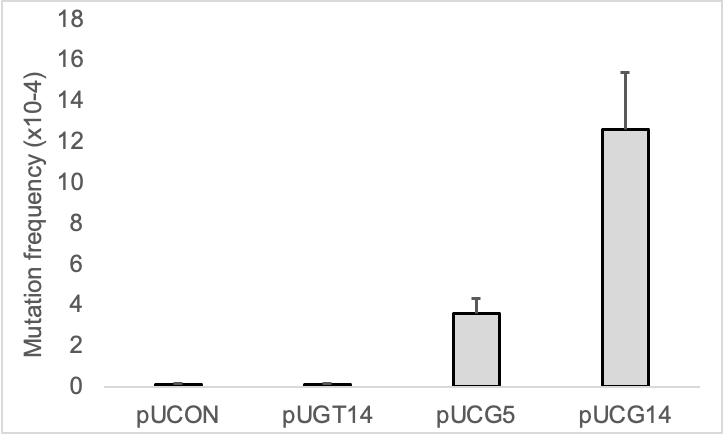

**Supplementary Figure 1: GT and CG repeat-induced mutation frequencies in DH5alpha cells.** GT14, CG5 and CG14 model sequences were cloned in a lacZ’ mutation reporter and the mutation frequencies were screened after the plasmids were replicated in DH5alpha cells for 16 hours. pUCON contains a random 28-bp sequence as a control. Error bars show the standard error of the mean from >3 independent repeats.

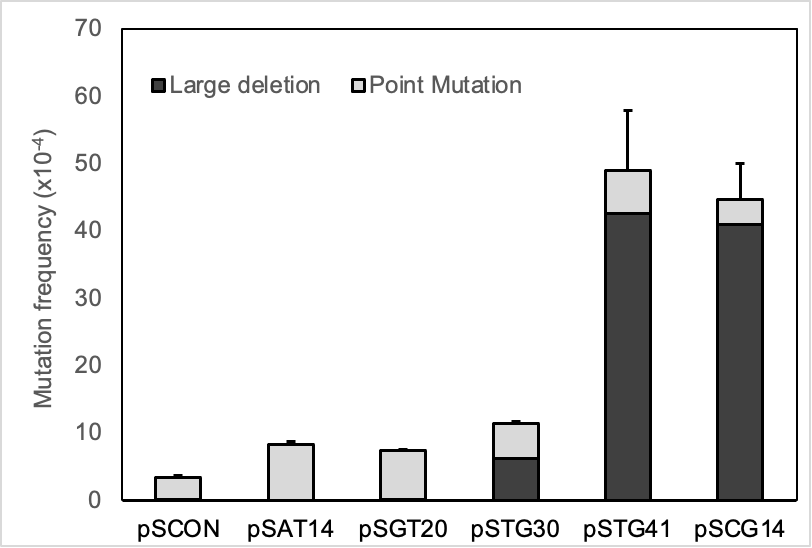

**Supplementary Figure 2: Repeat-induced mutation frequencies in mammalian COS-7 cells.** AT14, GT20, GT30, GT41 and CG14 model sequences were cloned in a supF mutation reporter and the mutation frequencies were screened after the mutation-reporter plasmids were transfected and replicated in COS-7 cells for 48 hours. pSCON contains a random 28-bp sequence as a control. >20 mutants were randomly picked and sequenced to determine the mutation spectra. Dark bars represent the frequencies of Z-DNA-induced large deletions resulting from DSBs, and light gray bars represent the frequencies of point mutations and small indels. Error bars show the standard error of the mean from >3 independent repeats. Results obtained for GT30 and GT41 were published as part of figure in (Xie et al. 2019).
